# Alpha oscillations in the human brain implement distractor suppression independent of target selection

**DOI:** 10.1101/721019

**Authors:** Malte Wöstmann, Mohsen Alavash, Jonas Obleser

**Affiliations:** Department of Psychology, University of Lübeck, Lübeck, Germany

**Keywords:** attention, selection, suppression, alpha oscillations, auditory

## Abstract

In principle, selective attention is the net result of target selection and distractor suppression. The way in which both mechanisms are implemented neurally has remained contested. Neural oscillatory power in the alpha frequency band (~10 Hz) has been implicated in the selection of to-be-attended targets, but there is lack of empirical evidence for its involvement in the suppression of to-be-ignored distractors. Here, we use electroencephalography (EEG) recordings of N = 33 human participants (males and females) to test the pre-registered hypothesis that alpha power directly relates to distractor suppression and thus operates independently from target selection. In an auditory spatial pitch discrimination task, we modulated the location (left vs right) of either a target or a distractor tone sequence, while fixing the other in the front. When the distractor was fixed in the front, alpha power relatively decreased contralaterally to the target and increased ipsilaterally. Most importantly, when the target was fixed in the front, alpha lateralization reversed in direction for the suppression of distractors on the left versus right. These data show that target-selection–independent alpha power modulation is involved in distractor suppression. While both lateralized alpha responses for selection and for suppression proved reliable, they were uncorrelated and distractor-related alpha power emerged from more anterior, frontal cortical regions. Lending functional significance to suppression-related alpha oscillations, alpha lateralization at the individual, single-trial level was predictive of behavioral accuracy. These results fuel a renewed look at neurobiological accounts of selection-independent suppressive filtering in attention.

**Significance statement:** Although well-established models of attention rest on the assumption that irrelevant sensory information is filtered out, the neural implementation of such a filter mechanism is unclear. Using an auditory attention task that decouples target selection from distractor suppression, we demonstrate that two sign-reversed lateralized alpha responses reflect target selection versus distractor suppression. Critically, these alpha responses are reliable, independent of each other, and generated in more anterior, frontal regions for suppression versus selection. Prediction of single-trial task performance from alpha modulation after stimulus onset agrees with the view that alpha modulation bears direct functional relevance as a neural implementation of attention. Results demonstrate that the neurobiological foundation of attention implies a selection-independent alpha oscillatory mechanism to suppress distraction.

## Introduction

Human goal-oriented behavior requires both, the selection of relevant target information and the suppression of irrelevant distraction. Although foundational theories of attention implied some form of distractor suppression (e.g., Broadbent, 1958; Treisman, 1964), different neural implementations of suppression are conceivable. On the one hand, suppression might be contingent on selection, meaning that distractors outside the focus of attention are suppressed automatically (Noonan, Crittenden, Jensen, & Stokes, 2018). On the other hand, distractor suppression might be an independent neuro-cognitive process (Aron, 2007) that adapts to changing characteristics of the distractor even in case the focus of attention is unchanged.

The power of brain oscillations in the alpha frequency band (~10 Hz) robustly tracks when humans shift their focus of attention between sensory modalities (Adrian, 1944; de Pesters et al., 2016; Fu et al., 2001), to time points of anticipated target presentation (Payne, Guillory, & Sekuler, 2013; Rohenkohl & Nobre, 2011), or to a particular location in space (Popov, Gips, Kastner, & Jensen, 2019; Sauseng et al., 2005; Worden, Foxe, Wang, & Simpson, 2000). Since alpha power drops in brain regions related to processing upcoming target stimuli (de Pesters et al., 2016), and since lower alpha power correlates with increased neural responses to the target (Gould, Rushworth, & Nobre, 2011; Wöstmann, Waschke, & Obleser, 2019) and enhanced behavioral measures of target detection (Thut, Nietzel, Brandt, & Pascual-Leone, 2006), low alpha power is considered a signature of enhanced neural excitability to support target selection.

At the same time, alpha power does increase in brain regions that process distracting stimuli. Although high alpha power is considered a brain state of inhibited neural processing (Foxe & Snyder, 2011; Jensen & Mazaheri, 2010; Strauß, Wöstmann, & Obleser, 2014), evidenced also by negative correlation of alpha power and brain activity measured in functional magnetic resonance imaging (fMRI; Laufs et al., 2003), it is unclear at present whether high alpha power constitutes an independent signature of distractor suppression or a by-product of target selection (Foster & Awh, 2018; Van Diepen, Foxe, & Mazaheri, 2019).

To study the contribution of alpha oscillations to attentional selection and suppression, neuroscientists have used spatial cueing of an upcoming target location under competing distraction at another location. Across modalities, target cueing induces alpha power lateralization, that is, alpha power decreases in the hemisphere contralateral to the target and increases in the ipsilateral hemisphere (Ahveninen, Huang, Belliveau, Chang, & Hamalainen, 2013; Bauer, Kennett, & Driver, 2012; Haegens, Handel, & Jensen, 2011). Alpha lateralization bears behavioral relevance: It is modulated stronger for correct than incorrect responses to the target stimulus (Haegens, Handel, et al., 2011; Wöstmann, Herrmann, Maess, & Obleser, 2016); and it modulates behavioral responses to the target if participants’ endogenous alpha lateralization is stimulated transcranially (magnetically: Romei, Gross, & Thut, 2010; or electrically: Wöstmann, Vosskuhl, Obleser, & Herrmann, 2018).

Critically, previous studies often confounded target and distractor location by design (e.g., Kelly, Lalor, Reilly, & Foxe, 2006; van Diepen, Miller, Mazaheri, & Geng, 2016; Wöstmann et al., 2016): Whenever the target appeared on the left, the distractor was presented on the right, and vice versa. To unambiguously assign alpha lateralization to selection versus suppression, it is necessary to physically decouple target and distractor location during spatial attention. Addressing this, we here disentangled this conundrum by fixing the position of either an auditory target or distractor stimulus in the front of the listener. We then only varied the respective other stimulus to come from either the left or right side.

We find that lateralized alpha power is an autonomous (i.e., target-independent) signature of suppression that can track the location of the distractor. The lateralization of alpha power proves as a reliable neural signature, separating selection from suppression within individuals and in their underlying neural generators. Finally, the instantaneous degree of alpha lateralization after, but not prior to, stimulus onset predicts trial-by-trial variations in behavioral accuracy for detecting small pitch changes in the target sound.

## Materials and Methods

### Pre-registration

Prior to data recording, we pre-registered the study design, data sampling plan, hypotheses, and analyses procedures online with the Open Science Framework (OSF; https://osf.io/bv7zs). All data and analysis code will be made available upon reasonable request.

### Participants

We analyzed data of N = 33 right-handed participants (M_age_ = 23.3 years; SD_age_ = 3.9 years; 22 females). Data of three additional participants were recorded but excluded, as two of them had excessive EEG artifacts and one was unable to perform the task. Participants were financially compensated or received course credit. All procedures were approved by the local ethics committee of the University of Lübeck.

### Auditory stimuli and task setup

Task design and stimuli were adapted from Dai, Best, and Shinn-Cunningham (2018). Due to a change in the stimulus sampling frequency, however, duration and pitch of auditory stimuli deviated slightly in the present study. The experiment was implemented in the Psychtoolbox (Brainard, 1997) for Matlab, and conducted in a soundproof cabin. All auditory stimuli were presented at a sampling frequency of 48 kHz at a comfortable level of ~65 dBA.

On each trial, an auditory spatial cue (10.9-kHz low-pass filtered Gaussian noise; 0.46 s) was presented at one location followed by two concurrent tone sequences presented at two different spatial locations (front, left, or right). Each tone sequence consisted of two 0.46-s complex tones (fixed ISI of 46 ms), one low-pitch tone and one high-pitch tone. All tones and the spatial cue were gated on and off with 92-ms cosine ramps.

The pitch of the low-pitch tone was fixed at 192.7 Hz (including 32 harmonics) for one sequence and at 300.4 Hz (including 2 harmonics) for the other sequence. Throughout the experiment, the fundamental frequency of the high-pitch tone in each sequence varied in semitones relative to the low-pitch tone using an adaptive tracking procedure (two-up-one-down) to arrive at ~71 % task accuracy (Levitt, 1971). Thus, after one incorrect response or two subsequent correct responses the pitch difference within each tone sequence was increased or decreased in steps of 0.05 semitones on the next trial, respectively. The initial pitch difference for the tracking procedure was obtained from a pre-experiment training session. The cue location (front vs side loudspeaker), the pitch direction within each sequence (increasing vs decreasing), and the assignment of tone sequences to the loudspeaker locations was balanced across trials and drawn randomly for an individual trial.

Tone sequences were presented in free field using a pair of loudspeakers (Logitech, x140). The location of a speaker could be either front or side (i.e., 0 or ±90 degrees azimuth relative to ear-nose-ear line). Loudspeakers were positioned at approximately 70 cm distance to the participant’s head. As the main experimental manipulation, the location of the side speaker changed between left and right across blocks of the experiment. To this end, the experimenter moved the side speaker accordingly in the breaks between blocks of the experiment. The other speaker was positioned in the front of the participant throughout. There were three blocks of the experiment with the side speaker positioned on the left side, and three blocks with the side speaker on the right. The order of blocks was counter-balanced across participants, and alternated between blocks with the side speaker on the left and right. Within each block a participant completed 96 trials (each loudspeaker served as the target in 48 trials).

### Experimental design

The present experimental design implemented four conditions. In experimental blocks with the distractor fixed in the front, the target was either on the left side (select-left condition) or on the right (select-right condition). In blocks with the target fixed in the front, the distractor was either on the left side (suppress-left condition) or on the right (suppress-right condition).

### Procedure

At the start of each trial, after a jittered period of ~1 sec (0.8–1.2 s), an auditory spatial cue was presented on one loudspeaker to inform the participant about the target loudspeaker location. After a jittered period of ~1.8 s (right-skewed distribution; median: 1.84 s; truncated at 1.47 and 2.48 s) relative to cue offset, two tone sequences were presented concurrently. Participants reported whether the tone sequence at the target location increased or decreased in pitch and how confident they were in this response using a response box with four buttons. Participants were instructed to fixate a cross in the middle of the response box throughout the experiment. Prior to the main experiment, a short training ensured that participants could perform the pitch discrimination task.

### EEG recording and preprocessing

The EEG was recorded at 64 active scalp electrodes (Ag/Ag-Cl; ActiChamp, Brain Products, München, Germany) at a sampling rate of 1000 Hz, with a DC–280 Hz bandwidth, against a left mastoid reference (channel TP9). All electrode impedances were kept below ~30 kOhm. To ensure equivalent placement of the EEG cap, the vertex electrode (Cz) was placed at 50% of the distance between inion and nasion and between left and right ear lobes.

For EEG data analysis, we used the FieldTrip toolbox (Oostenveld, Fries, Maris, & Schoffelen, 2011) for Matlab (R2013b/R2018a) and custom scripts. Offline, the continuous EEG data were filtered (1-Hz high-pass; 100-Hz low-pass) and segmented into epochs relative to the onset of the spatial cue (−2 to +6 s). An independent component analysis (ICA) was used to detect and reject components corresponding to eye blinks, saccadic eye movements, muscle activity, and heartbeat. On average 38.48 % (SD: 10 %) of components were rejected. After visual inspection of EEG time-domain data, noisy electrodes (one electrode of two participants) were interpolated using the nearest neighbor approach implemented in FieldTrip. Finally, trials for which an individual EEG channel exceeded a range of 200 microvolts were rejected. On average 568 trials (SD: 11 trials) of executed 576 trials per participant were used for further analyses.

### Analysis of neural oscillatory activity

EEG data were re-referenced to the average of all electrodes and down-sampled to 250 Hz. Single-trial time-frequency representations were derived using complex Fourier coefficients for a moving time window (fixed length of 0.5 s; Hanning taper; moving in steps of 0.04 s) for frequencies 1–30 Hz with a resolution of 1 Hz.

To quantify the impact of selection and suppression, single-trial power representations (squared magnitude of complex Fourier coefficients) were calculated for individual experimental conditions: select-left, select-right, suppress-left, and suppress-right. For each participant, two lateralization indices (LI) were calculated on absolute oscillatory power (Pow). The first index quantifies oscillatory signatures of target selection:

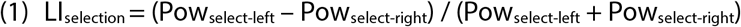

Importantly, the second index quantifies oscillatory signatures of distractor suppression, which goes beyond what previous studies have analyzed:

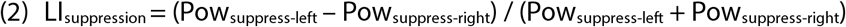

For statistical analyses, we followed our pre-registered analysis plan (https://osf.io/bv7zs). In brief, we averaged each lateralization index (LI) across frequencies in the alpha band (8–12 Hz), the time interval from cue onset to earliest tone sequence onset (0–1.9 s), separately for two sets of 12 left- and 12 right hemispheric occipito-parietal electrodes (TP9/10, TP7/8, CP5/6, CP3/4, CP1/2, P7/8, P5/6, P3/4, P1/2, PO7/8, PO3/4, and O1/2). For statistical comparisons of the LI (left vs. right hemisphere; selection vs. suppression), we used non-parametric permutation tests. The reported *p*-value corresponds to the relative number of absolute values of 10,000 dependent-samples *t*-statistics computed on data with permuted condition labels exceeding the absolute empirical *t*-value for the original data.

To determine reliability of lateralization indices, we divided each participant’s trials into three consecutive portions (each consisting of approximately 192 trials, with 48 trials for each condition), followed by calculation of lateralization indices for each portion. Next, we calculated the reliability metric Cronbach’s Alpha (CA) for each lateralization index across the three portions. The *p*-value for CA was derived by the relative number of permuted CAs, derived from 10,000 permutations of single-subject lateralization indices within each one of the three portions, exceeding the empirical CA (Prelog, Berry, & Mielke, 2009).

For the non-significant Spearman correlation of the two lateralization indices (LI_slection_, LI_suppression_) we report the Bayes Factor (computed for Kendall’s tau in the software Jamovi). The Bayes Factor (*BF*) indicates how many times more likely the observed data are under the alternative (H_1_) compared to the null hypothesis (H_0_). By convention, a *BF* > 3 begins to lend support to H_1_, whereas a *BF* < 0.33 begins to lend support to H_0_ (Dienes, 2014).

### Control for saccadic eye movements

Although participants were instructed to keep central gaze during the entire experiment, it might be that systematic differences in saccadic eye movements confounded the results (see e.g. Quax, Dijkstra, van Staveren, Bosch, & van Gerven, 2019), even in an auditory attention task. To rule this out, we inspected the EEG for independent components tuned to horizontal saccadic eye movements.

Prior to the main experiment, each participant performed a brief eye movement task. Participants followed a dot on the screen that jumped eight times either vertically (up and down by ~8° visual angle) or horizontally (left and right by ~8° visual angle) with inter-jump-intervals of 1s. The task started with a horizontal movement trial and then alternated between vertical and horizontal movement trials.

Trials of the eye movement task were segmented into epochs relative to the onset of the first jump of the dot (−1 to +10s). An independent component analysis (ICA) was used to extract one component for horizontal eye movements for each participant (vertical eye movements were not considered further in this study). The event-related potential (ERP) across all participant’s horizontal eye movement components clearly differentiated between saccades to the left versus right side (Fig. 1A).

**Figure 1.**
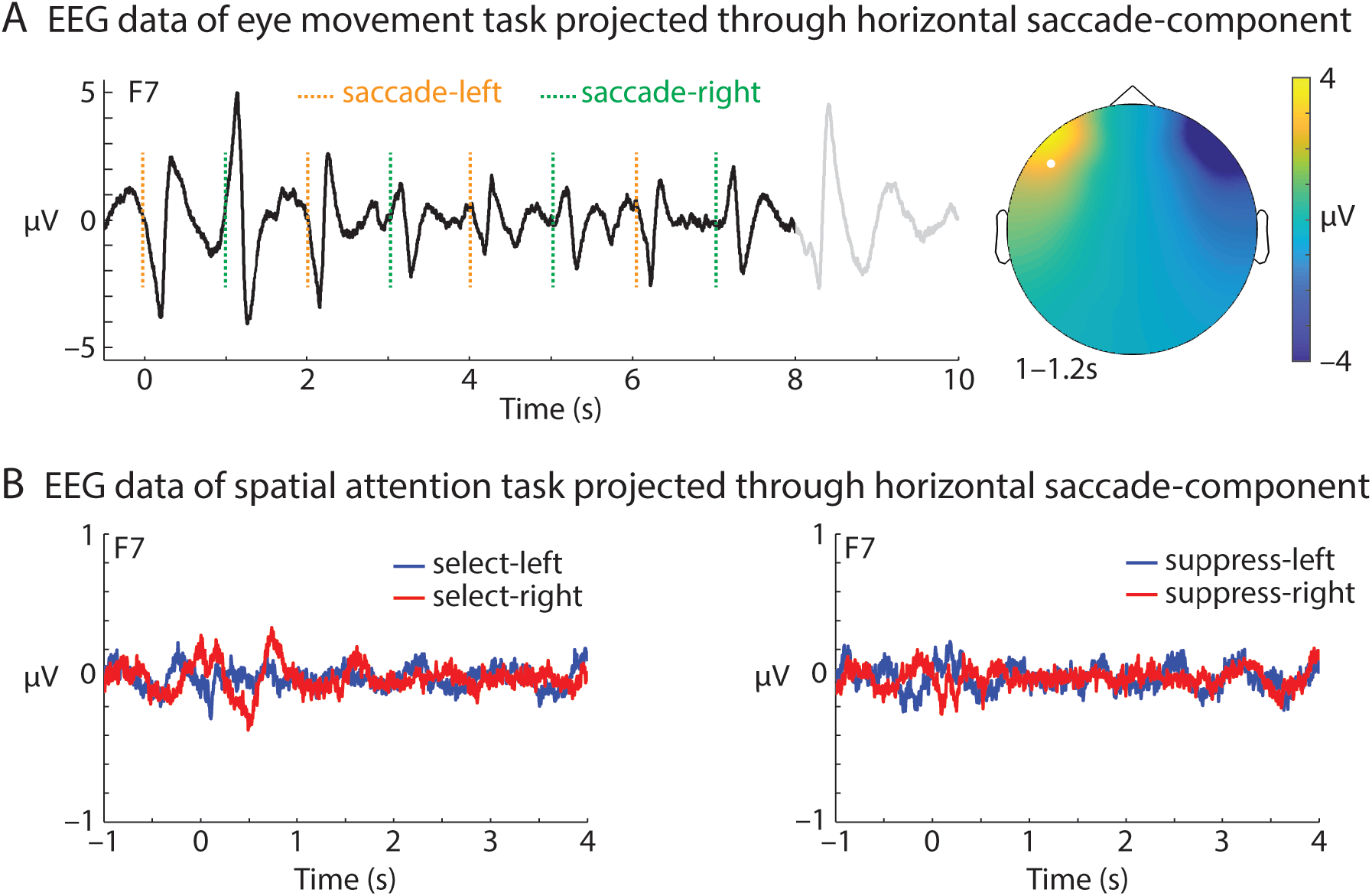
(**A**) During a pre-experiment eye movement task participants followed a dot, which jumped eight times from the left to the right side on the screen, with their gaze. The grand-average event-related potential (ERP; at electrode F7) was computed on the EEG data projected through one horizontal saccade component per participant (N = 33). At 8 s, participants performed an additional saccade back to the center of the screen to read task instructions. The topographic map shows ERP amplitude (1–1.2s) following the onset of one saccade to the right side (electrode F7 highlighted). (**B**) EEG data during the spatial attention task were projected through the same horizontal saccade component for each participant as in (A). Grand-average ERPs time locked to the onset of the auditory spatial cue show no obvious saccade-related activity in trials with targets or distractors on the left versus right side.

To control for potential confounds of horizontal saccadic eye movements in the EEG data of the spatial attention task, we projected each participant’s raw task data (with no trials rejected) through the horizontal eye movement component, followed by computation of the ERP. For statistical analysis, we performed two cluster-based permutation tests to contrast the ERP during trials of the spatial attention task (0 to 4 s) for target selection on the left versus right side and distractor suppression on the left versus right side, respectively. In essence, these cluster-based permutation tests cluster *t*-values of adjacent bins in time-electrode space (minimum cluster size: 3 adjacent electrodes) and compare the summed *t*-statistic of the observed cluster against 10,000 randomly drawn clusters from the same data with permuted condition labels. The *p*-value of a cluster corresponds to the proportion of Monte Carlo iterations in which the summed *t*-statistic of the observed cluster is exceeded (two-sided testing; alpha level of 0.05).

ERPs of task data projected through the horizontal eye movement component did not show differences between experimental conditions (Fig. 1B). Cluster permutation tests revealed no significant differences in the ERP for selection of targets on the left versus right side (all cluster *p*-values > 0.19) or suppression of distractors on the left versus right side (all cluster *p*-values > 0.36).

In an additional analysis (not shown), we also tested for gamma power (> 30 Hz) differences in task data projected through horizontal eye movement components, which might be indicative of microsaccadic eye movements (Yuval-Greenberg, Tomer, Keren, Nelken, & Deouell, 2008). Two cluster permutation tests on time-frequency representations of oscillatory power in a broad frequency range (1–90 Hz) revealed no significant power differences for select-left versus select-right or suppress-left versus suppress-right conditions (all cluster *p*-values > 0.2).

These results suggest that participants complied with our task instructions and did not systematically perform saccadic eye movements to or away from the to-be-selected or to-be-suppressed loudspeaker.

### EEG source analysis

We used the Dynamic Imaging of Coherent Sources (DICS) beamformer approach (Gross et al., 2001) implemented in FieldTrip. A standard head model (Boundary Element Method, BEM; 3-shell) was used to calculate leadfields for a grid of 1 cm resolution. Spatial filters were calculated from the leadfield and the cross-spectral density of Fourier transforms centered at 10 Hz with ± 2 Hz spectral smoothing in the time interval 0–1.9 s relative to cue onset. For each participant, two spatial filters were calculated to source-localize LI_selection_ and LI_suppression_, based on all trials with the target or distractor on the side, respectively. The spatial filter for LI_selection_ was used to localize alpha power separately for select-left and select-right trials, followed by calculation of the lateralization index LI_selection_ on the source level (and accordingly for LI_suppression_). Finally, source-level LIs were averaged across participants and mapped onto a standard brain surface.

### Combined behavioral and EEG data analysis

Single-trial EEG and behavioral data were matched. Due to few missing EEG triggers, one participant’s data were excluded from behavioral analyses. For remaining N = 32 participants, an average of 523 trials was used. Behavioral data were analyzed using mixed-effects models implemented in the *fitlme* function for Matlab. The response variable of interest was single-trial confidence-weighted accuracy, derived by transformation of binary accuracy into 1 and 1/3 for correct responses with respective high and low confidence, and into −1 and −1/3 for incorrect responses with respective high and low confidence (Wöstmann, Herrmann, Wilsch, & Obleser, 2015). As predictors, we used the titrated pitch difference (within both tone sequences), congruency of pitch direction across the two tone sequences (congruent versus incongruent), location of lateralized loudspeaker (left versus right), and role of lateralized loudspeaker (target versus distractor), with participant as a random intercept term, resulting in the linear-model expression: Confidence-weighted accuracy ~ 1 + Titrated pitch difference + Congruency of pitch direction × Location of lateralized loudspeaker × Role of lateralized loudspeaker + (1|Participant ID) To model the relation of alpha lateralization and confidence-weighted accuracy, we included single-trial alpha lateralization (LI_single-trial_) before tone sequence onset (0–1.9 s) and for a 1-s time window moving in 0.1-s steps through a trial as predictors in separate linear mixed-effects models. Single trial alpha power lateralization was quantified as the contrast of alpha power (obtained via Fourier transform using multi-tapering at 10 Hz with 2-Hz spectral smoothing) at 12 parieto-occipital left-minus-right hemispheric electrodes:

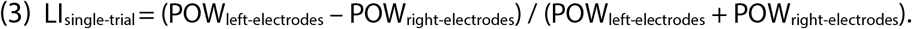

## Results

Participants (N = 33) performed a spatial pitch-discrimination task (Fig. 2A; adopted from Dai et al., 2018). The loudspeaker setup changed blockwise between front-and-left or front-and-right, with one of the two loudspeakers serving as target and the other as distractor on each trial. A cue tone in the beginning of each trial indicated the location of the target loudspeaker. Participants had to report whether the ensuing tone sequence at the target location increased or decreased in pitch. A distracting tone sequence was presented simultaneously by the other loudspeaker.

**Figure 2.**
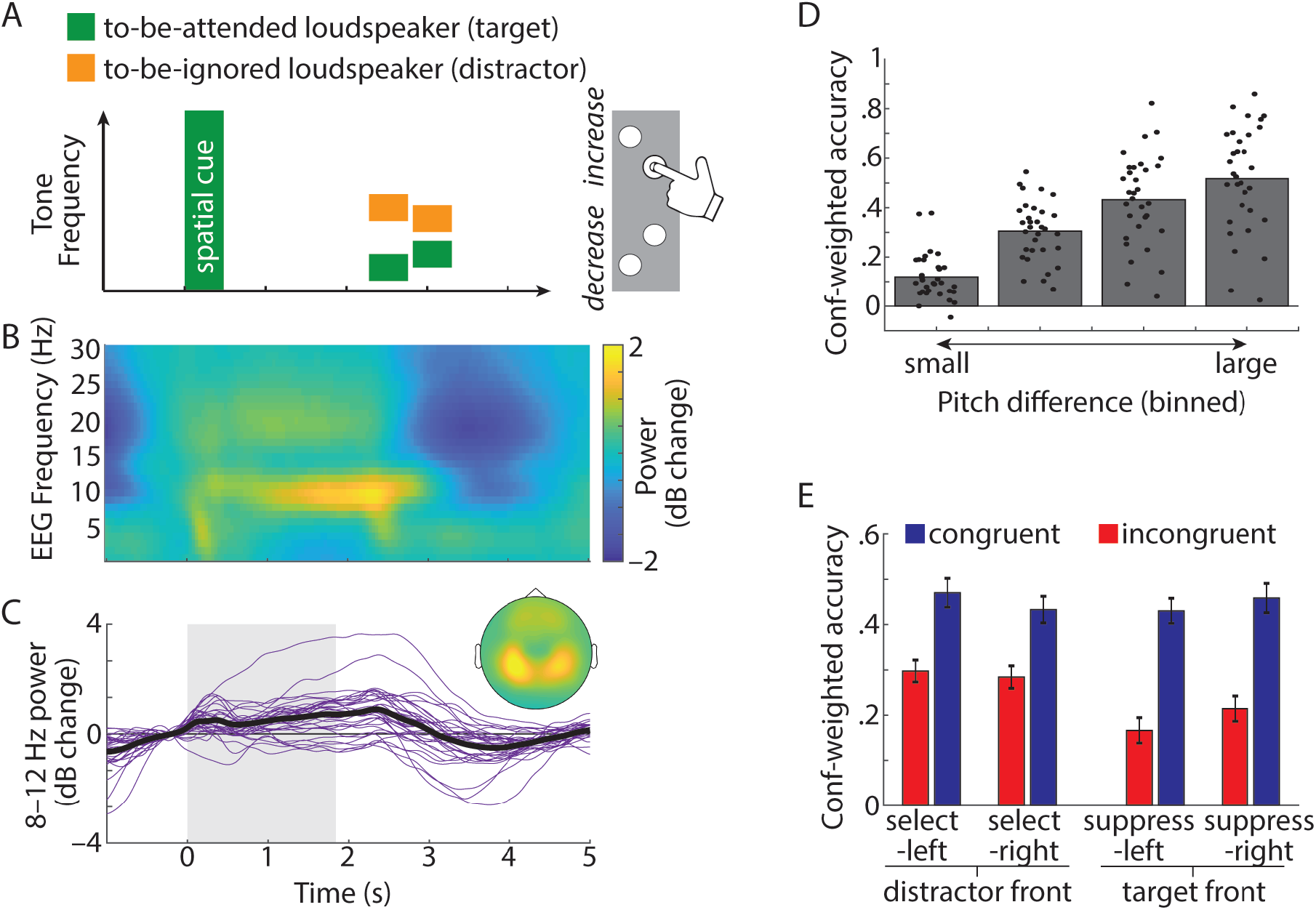
(**A**) Trial design. Presentation of a broad-band auditory cue (0–10.9 kHz; 0.46 s) was followed by a jittered silent interval (1.47–2.48 s). Next, two tone sequences, each consisting of two brief (0.46 s) complex tones, were presented at different locations (front, left, or right). Fundamental frequencies of low-frequency tones within each sequence were fixed at 193 and 300 Hz. Frequencies of high-frequency tones were titrated throughout the experiment. Participants had to judge whether the target tone sequence at the cued location had increased or decreased in pitch. Participants also indicated confidence in their judgement (high: outer buttons; low: inner buttons). (**B**) Time-frequency representation of oscillatory power (relative to a pre-trial baseline; −0.5 to 0 s; in decibel (dB) change), averaged across N = 33 participants and all (64) scalp electrodes. (**C**) Thin purple lines show single-subject alpha power (8–12 Hz) time courses; thick black line shows average; topographic map shows average 8–12 Hz power from 0 to 1.9 s (grey box). (**D**) Bars show average confidence-weighted accuracy as a function of the pitch difference between the two tones within each tone sequence (divided into four bins for each participant for visualization), which was titrated over the course of the experiment. Dots show single-subject data. (**E**) Bars and error bars show average ±1 between-subject SEM of confidence-weighted accuracy, separately for congruent trials (both tone sequences increasing/decreasing in pitch; blue) and incongruent trials (one tone sequences increasing and other decreasing in pitch; red).

As commonly observed in auditory attention tasks (e.g. Henry, Herrmann, Kunke, & Obleser, 2017; Wöstmann, Fiedler, & Obleser, 2017; Wöstmann et al., 2015), power in the alpha frequency band (8-12 Hz) at parietal electrodes relatively increased after trial onset and decreased in the end of a trial before participants performed a behavioral response (Fig. 2 B&C).

### Behavioral results

To avoid ceiling- and floor performance, the pitch difference within both tone sequences was titrated throughout the experiment (average pitch difference = 0.339 semitones; between-subject SD = 0.524), using an adaptive procedure to target a proportion of ~0.71 correct responses. Average proportion correct was 0.715 (between-subject SD: 0.044) and average response time was 0.955 s (between-subject SD: 0.523). We modeled single-trial confidence-weighted accuracy (see Fig. 2-2 and 2-3 for analysis of raw accuracy and response time) on predictors *titrated pitch difference* (within both tone sequences), *congruency of pitch direction across target and distractor tone sequence* (congruent versus incongruent), *location of lateralized loudspeaker* (left versus right), and *role of lateralized loudspeaker* (target versus distractor).

Confidence-weighted accuracy increased if the pitch difference between the tones within each sequence (target and distractor) was larger (Fig. 2D; *t_16949_* = 10.031; *p* < 0.001), if tone sequences were congruent in pitch direction (Fig. 2E *t_16949_* = 8.572; *p* < 0.001), and if the target loudspeaker was on the side compared to front (main effect role of lateralized loudspeaker: *t_16949_* = −5.622; *p* < 0.001), which agrees with a previous study that used a similar task design (Dai et al., 2018). The latter effect was driven by incongruent trials, evidenced by the congruency × role of lateralized loudspeaker interaction (*t_16949_* = 2.764; *p* = 0.006). The location × role of lateralized loudspeaker interaction approached statistical significance (*t_16949_* = 1.803; *p* = 0.071), which speaks to a tendency of higher confidence-weighted accuracy for target selection on the left versus right but for distractor suppression on the right versus left (see Fig. 2-1, 2-2, and 2-3 for complete summary tables of mixed model analyses).

### Alpha lateralization tracks target location independent of distractor

We tested whether alpha power would track the spatial location of the to-be-selected target stimulus (left vs. right) under fixed distraction from the front. To this end, we contrasted alpha power during the anticipation of tone sequences (0–1.9 s) for trials with a target on the left versus right side (Fig. 3 A&B). Alpha power relatively increased in parietal, occipital, and inferior temporal regions in the hemisphere ipsilateral to the cued target location, and decreased contralaterally. Accordingly, the lateralization index (LI_selection_), which quantifies the difference in lateralized alpha power at left-minus-right parieto-occipital electrodes, was significantly positive (*CI_95_* = [0.033; 0.06]; *permutation p-value* < 0.001; see Fig. 3-1 for time- and frequency-resolved lateralization indices).

**Figure 3.**
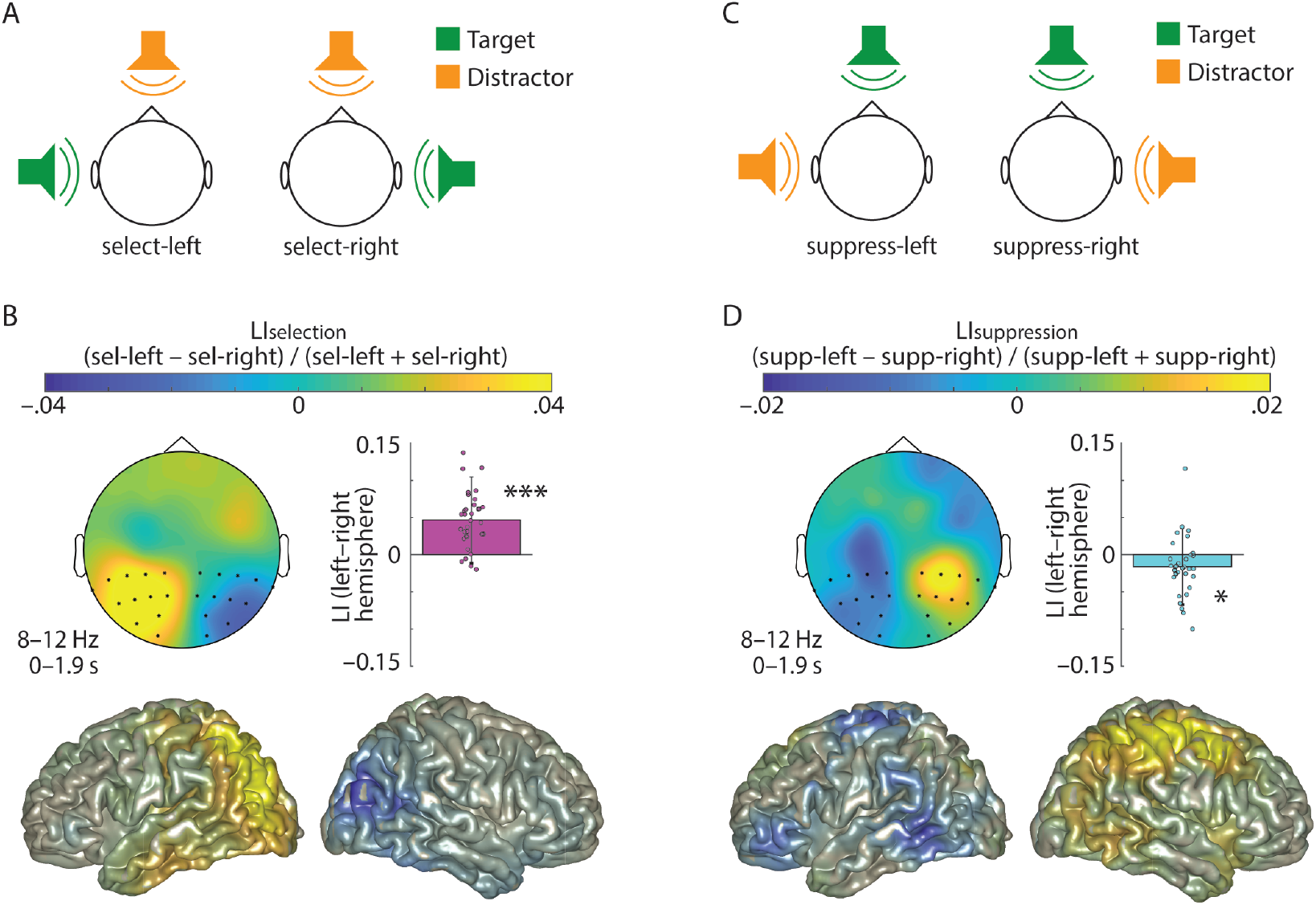
(**A**) Schematic illustration of task setup in select-left and select-right trials. (**B**) Topographic map and cortical surfaces show the lateralization index to contrast select-left and select-right trials under fixed distraction from the front (LI_selection_). The lateralization index was calculated for 8–12 Hz alpha power during anticipation of tone sequences (0–1.9 s). Bar graph, error bar, and dots show average, ±1 between-subject SEM, and single-subject differences of LI for highlighted parieto-occipital electrodes on the left versus right hemisphere, respectively. (**C&D**) Same as A&B, but for the lateralization index for suppression of lateralized distractor stimuli (LI_suppression_). * p < 0.05; *** p < 0.001.

### Alpha lateralization tracks distractor location independent of target

The most important objective of the present study was to test whether lateralized alpha power contains also a neural signature of distractor suppression, independent of target selection. To test this, we contrasted alpha power for trials with the target stimulus fixed in the front but with an anticipated distractor on the left versus right side (Fig. 3 C&D). As predicted, the location of a to-be-suppressed distractor modulated alpha power orthogonally to target selection: Alpha power relatively increased in parietal, posterior temporal, and frontal lobe regions in the hemisphere contralateral to the distractor and decreased ipsilaterally. Accordingly, the lateralization index (LI_suppression_) at parieto-occipital electrodes on the left versus right hemisphere was significantly negative (*CI_95_* = [−0.03; − 0.002]; *permutation p-value* = 0.0202).

Having established EEG signatures of target selection (LI_selection_) and distractor suppression (LI_suppression_), we tested reliability of these signatures. Only if they are sufficiently reliable, meaning that results on repeated tests correlate positively, any relation or difference between the two signatures can be interpreted in a meaningful way. We divided each participant’s data into three consecutive portions and found significant positive values of the reliability metric Cronbach’s Alpha (CA) of alpha lateralization for target selection (LI_selection_; *CA* = 0.602; *permutation p-value* = 0.001) and distractor suppression (LI_suppression_; *CA* = 0.522; *permutation p-value* = 0.006).

### Alpha signatures of selection and suppression operate independently

As for a potential hierarchy between alpha mechanisms of selection versus suppression, we tested whether anticipation of to-be-selected targets induces stronger alpha lateralization than anticipation of to-be-suppressed distractors. The data speak clearly in favor of this hypothesis: The hemispheric difference in alpha lateralization (LI; bar graphs in Fig. 3) was significantly more positive for LI_selection_ than it was negative for LI_suppression_ (*CI_95_* = [0.008; 0.053]; *permutation p-value* = 0.005).

We then asked, to what extent alpha lateralization for target selection and distractor suppression would relate. If the two neural signatures instantiate the same underlying cognitive faculty, participants with stronger alpha lateralization for target selection should show stronger alpha lateralization for distractor suppression as well. This would result in more positive LI_selection_ being accompanied with more negative LI_suppression_ and thus in a negative correlation. To the contrary, LI_selection_ and LI_suppression_ were not significantly correlated (Fig. 4A; *r_Spearman_* = 0.153; *p* = 0.393; *BF* = 0.294). Thus, alpha lateralization for target selection and distractor suppression can be considered two largely independent neural signatures.

**Figure 4.**
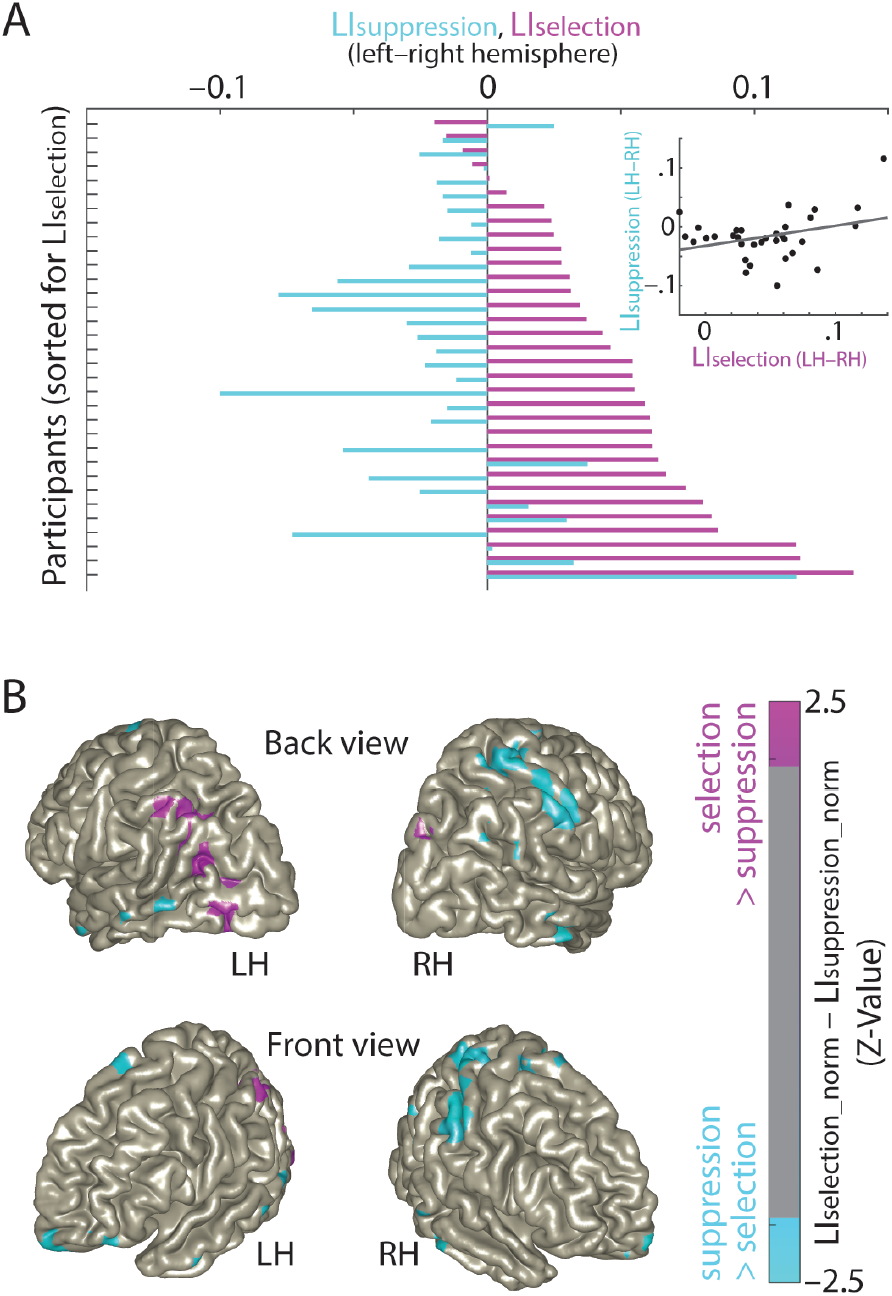
(**A**) Bars show single-subject alpha lateralization indices at left minus right parieto-occipital electrodes (pink: LI_selection_; blue: LI_suppression_), sorted for LI_selection_. Inset shows scatterplot for LI_selection_ and LI_suppression_, with least-squares regression line (*Spearman rho* = 0.153; *p* = 0.393). (**B**) Normalization of lateralization indices on the source-level was accomplished by z-transformation of single-subject lateralization indices, followed by taking the magnitude. Brain surfaces show Z-statistics derived from dependent-samples t-tests for the contrast of normalized lateralization indices: LI_selection_norm_ versus LI_supression_norm_. Z-statistics on brain surfaces are masked in case |Z| < 1.96, corresponding to p > 0.05 for two-sided testing (uncorrected). Top: back view; bottom: front view. LH: left hemisphere; RH: right hemisphere.

Next, we tested whether alpha lateralization for target selection and distractor suppression might be implemented by different neural generators. To this end, we focused on differences in spatial distribution but not strength or direction of the hemispheric difference of alpha power modulation. We z-transformed each participant’s lateralization indices, followed by taking their magnitudes (referred to as LI_selection_norm_ and LI_suppression_norm_). Indeed, neural generators of selection and suppression differed significantly: Target selection was driven by relatively stronger alpha power modulation in parieto-occipital cortex regions, primarily in the left hemisphere (pink regions in Fig. 4B). Conversely, neural generators of distractor suppression were modulated relatively stronger in more widespread, right superior and inferior parietal, inferior temporal, superior frontal, and middle frontal cortex regions (blue regions in Fig. 4B).

### Single-trial alpha lateralization predicts task performance

To test our hypothesis that lateralized alpha power predicts task performance, we added single-trial alpha lateralization (LIsingle-trial) before the onset of tone sequences (0–1.9 s) to the set of predictors used in the analysis of behavioral results (Fig. 2 D&E). Relatively higher left-than-right hemispheric alpha power should be beneficial in select-left and suppress-right trials but detrimental in select-right and suppress-left trials, reflected in a location × role of lateralized loudspeaker × alpha lateralization interaction. Against what we had hypothesized, this interaction was clearly non-significant (*t_16941_* = 0.002; *p* = 0.999). However, when we used single-trial alpha lateralization for successive time intervals throughout a trial as predictors (in separate mixed-effects models) in an exploratory follow-up analysis, we found a weak but significant location × role of lateralized loudspeaker × alpha lateralization interaction only in a late time window in the end of tone sequence presentation (3.1–3.4 s; Fig. 5A). The interaction was driven by trials with the lateralized loudspeaker on the right side (Fig. 5B): In line with the proposed inhibitory role of alpha power, relatively higher left-hemispheric alpha power improved suppression of distractors on the right but impaired selection of targets on the right.

**Figure 5.**
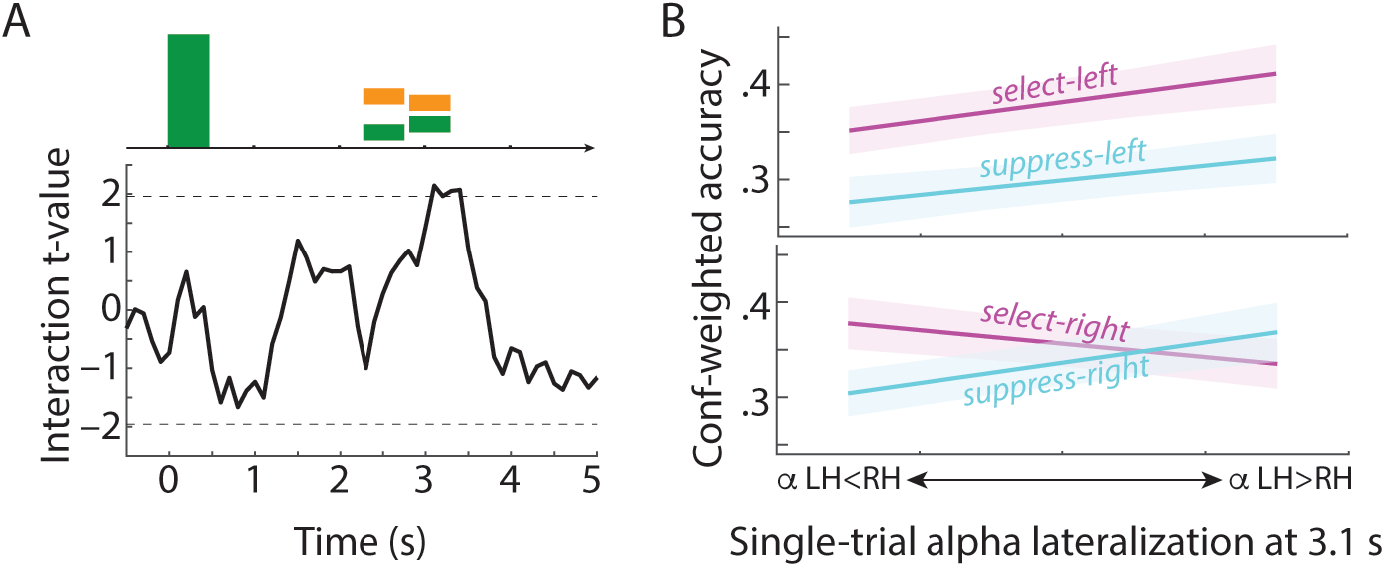
(**A**) Black solid line shows trace of t-values for the location × role of lateralized loudspeaker × single-trial alpha lateralization interaction obtained from multiple linear mixed models. Dashed lines indicate significance thresholds (*alpha* = 0.05; uncorrected). (**B**) Visualization of significant location × role of lateralized loudspeaker × alpha lateralization interaction at a latency of 3.1 s. For each participant and experimental condition, single-trial alpha lateralization values (LI_single-trial_; quantifying left-versus-right hemispheric alpha power) were separated into four bins, followed by fitting a linear slope to average confidence-weighted accuracy as a function of increasing bin number. Lines show average across single-subject linear fits, shaded areas show ±1 between-subject SEM. LH: left hemisphere; RH: right hemisphere.

## Discussion

The present study tested whether the human brain implements a mechanism of distractor suppression that is independent of target selection. Results show that this is the case. Under fixed presentation of auditory targets in the front, anticipation of to-be-suppressed distractors on the left versus right side induced contralateral increase and ipsilateral decrease of alpha power. Alpha lateralization for target selection and distractor suppression were not only opposite in direction but we demonstrate here that the two are reliable and independent neural signatures generated in partly distinct underlying neural networks. Exploratory analysis of the relation between alpha lateralization and behavioral task performance supports the view that alpha power modulation following stimulus onset controls the read-out of sensory objects that compete for attention.

### Alpha lateralization is an autonomous signature of distractor suppression

The most important result of the present study was significant lateralization of alpha power in anticipation of distractors on the left versus right side in case the focus of attention was fixed to the front (Fig. 3C&D). This finding confirms our pre-registered hypothesis of alpha lateralization as a neural signature of target-independent (i.e., autonomous) distractor suppression (https://osf.io/bv7zs).

While neuroscience has previously shown that alpha oscillations represent an inhibitory signal that correlates with suppressed neural firing (Haegens, Nacher, Luna, Romo, & Jensen, 2011; Jensen, Bonnefond, & VanRullen, 2012), we demonstrate here that alpha power constitutes inhibition also in a psychological sense (Aron, 2007): It signifies a participant’s intention to ignore distracting sensory information. Our results clearly challenge the view that alpha oscillations are not involved in distractor suppression (Foster & Awh, 2018) and instead show that lateralized alpha power adapts to the location of anticipated distraction independent of selecting a target at a fixed location.

Participants in the present study made use of alpha power lateralization, although in opposite direction, to implement target selection versus distractor suppression. In this sense, alpha power is a neural signal that implements the psychological construct of an attentional filter (Lakatos et al., 2013; Wöstmann et al., 2016) to coordinate the interplay of selection and suppression. Since alpha responses for selection and suppression were reliable and uncorrelated, we consider these neural responses traits that allow to place a participant’s instantiation of the attentional filter in a two-dimensional space defined by neural target selection and distractor suppression (Fig. 4).

In theory, separation of competing target and distractor can be achieved by target enhancement or by distractor inhibition. So why should the human brain implement both of these mechanisms if one would suffice? For any biological system, the magnitude of response has a limited dynamic range. Attentional selection can only enhance neural processing of the target to a finite extent. Thus, a dual mechanism that additionally implements distractor suppression is able to effectively double the separation of target and distractor (Houghton & Tipper, 1984).

Neural implementation of target selection and distractor suppression. In line with the prevalent model of alpha power as a signature of neural inhibition (Foxe & Snyder, 2011; Jensen & Mazaheri, 2010; Strauß et al., 2014), relatively high alpha power in the hemisphere contralateral to a distractor indicates suppression, while low alpha power contralateral to a target indicates selection. Previous studies, which tied spatial locations of target and distractor by presenting either one left and the other right, likely observed a superposition of two underlying lateralized alpha responses that we separated in the present study.

Evidence for the involvement of alpha oscillations in neural processing of distraction comes also from studies that found alpha power modulation associated with changing features of the distractor, such as its acoustic quality (Wöstmann, Lim, & Obleser, 2017), visual similarity to the target (Bonnefond & Jensen, 2012), the number of distractors (Sauseng et al., 2009), or continuous luminance changes in the distractor (Jia, Fang, & Luo, 2019). The design of the present study allowed us to trace and contrast the neural sources of lateralized alpha responses for target selection and distractor suppression, which were prominent in superior and inferior parietal cortex region that are part of the dorsal attention network (Sadaghiani et al., 2010) and involved in coding spatial locations of stimuli in the environment (Colby & Goldberg, 1999).

The apparent absence of alpha lateralization in auditory cortex regions is not unusual for auditory spatial attention tasks in the EEG (Tune, Wöstmann, & Obleser, 2018). Attentional modulation of alpha power in auditory cortex regions has been demonstrated mainly using brain imaging methods with high spatial specificity such as Magnetoencephalography (Wöstmann et al., 2016) or intracranial recordings (Gomez-Ramirez et al., 2011). Since the net EEG signal is dominated by widespread parieto-occipital alpha power, sophisticated analyses procedures such as the subtraction of visually- and haptically-induced alpha modulation from auditory-induced alpha modulation (Spitzer, Fleck, & Blankenburg, 2014) are necessary to reveal dominant sources of alpha modulation in auditory cortex regions. Furthermore, similar patterns of EEG alpha power lateralization in parietal cortex regions have been found for spatial shifts of visual but also auditory attention (Banerjee, Snyder, Molholm, & Foxe, 2011), suggesting that parietal alpha modulation is a signature of supramodal spatial attention. In this respect, it is likely that parietal alpha power lateralization in the present study reflects the allocation of supramodal attention rather than unimodal auditory attention alone.

Compared to target selection, distractor suppression induced relatively stronger alpha modulation in distributed regions including parietal, and (pre-) frontal cortex (Fig. 4). Such a pattern of frontal and parietal activations, termed multiple-demand (MD) system, has been found involved in a multitude of cognitively challenging tasks (Duncan, 2010). In particular, prefrontal cortex is a source of executive control (Miller & Cohen, 2001; Sadaghiani & Kleinschmidt, 2016), which is crucial for the orchestration of different neural processes to implement attention. Prefrontal cortex involvement in distractor suppression has also been evidenced by its relation to performance in tasks requiring cognitive control, such as the Stroop task (e.g., Vendrell et al., 1995). Our results thus help integrate the lateralized neural alpha response to distraction into models of prefrontal cortex involvement in inhibitory cognitive control.

It is of note that direct comparison of target selection and distractor suppression is somewhat limited in the present study. A bottom-up auditory cue drew attention to the target, whereas participants had to infer that the distractor would occur at the non-cued location. Furthermore, it might in theory be possible that lateralized alpha oscillations code the relative position of the target with respect to the distractor, which was the same (i.e. 90° to the left) in select-left and suppress-right trials. However, our results clearly speak against this, since alpha lateralization for target selection and distractor suppression were not only different in strength and source origin, but were also uncorrelated.

### Behavioral relevance of the lateralized alpha response

Against what we had hypothesized, we did not find a significant relation of single-trial alpha lateralization prior to stimulus presentation and task performance (for a study that recently reported such a relation in a different task setting, see Dahl, Ilg, Li, Passow, & Werkle-Bergner, 2019). In previous studies, we observed that the strength of the relation of alpha power modulation and behavior varied considerably between task settings, with some studies finding robust relations (Wöstmann et al., 2016; Wöstmann et al., 2015), while other found weak (Wöstmann, Schmitt, & Obleser, in press) or no such relation (Wöstmann, Lim, et al., 2017).

In an exploratory follow-up analysis we found that alpha lateralization after the onset of competing tone sequences predicted single-trial pitch discrimination performance, although the relation was relatively weak. There is an ongoing debate whether alpha lateralization is a purely proactive mechanism of attentional control to prepare for upcoming target and distractor, or whether alpha lateralization also has the potency to reactively select the target and to suppresses the distractor after these have been encoded (for review of proactive and reactive mechanisms, see Geng, 2014).

Our results differentiate proactive and reactive accounts of attention: The observed alpha power lateralization prior to competing tone sequences speaks to alpha lateralization as a signature of proactive attentional control. However, prediction of pitch discrimination performance only by post-stimulus alpha lateralization signifies the behavioral relevance of lateralized alpha power as a mechanism of reactive attentional control (Wöstmann et al., 2016).

## Conclusion

Although well-established models of attention rest on the assumption that irrelevant sensory information is filtered out, the neural implementation of such a filter mechanism is unclear. Using a task design that decouples target selection from distractor suppression, we demonstrate here that two independent lateralized alpha responses reflect target selection versus distractor suppression. These alpha responses are sign-reversed, reliable, and originate in more frontal, executive cortical regions for suppression than selection. Furthermore, lateralized alpha power predicts participants’ accuracy in the judgement of a pitch change in the target stimulus. These findings support so-called “active suppression” models of attention, in which suppression is not a necessary by-product of selection but an independent neuro-cognitive process.

## Extended data

**Figure 2-1.**
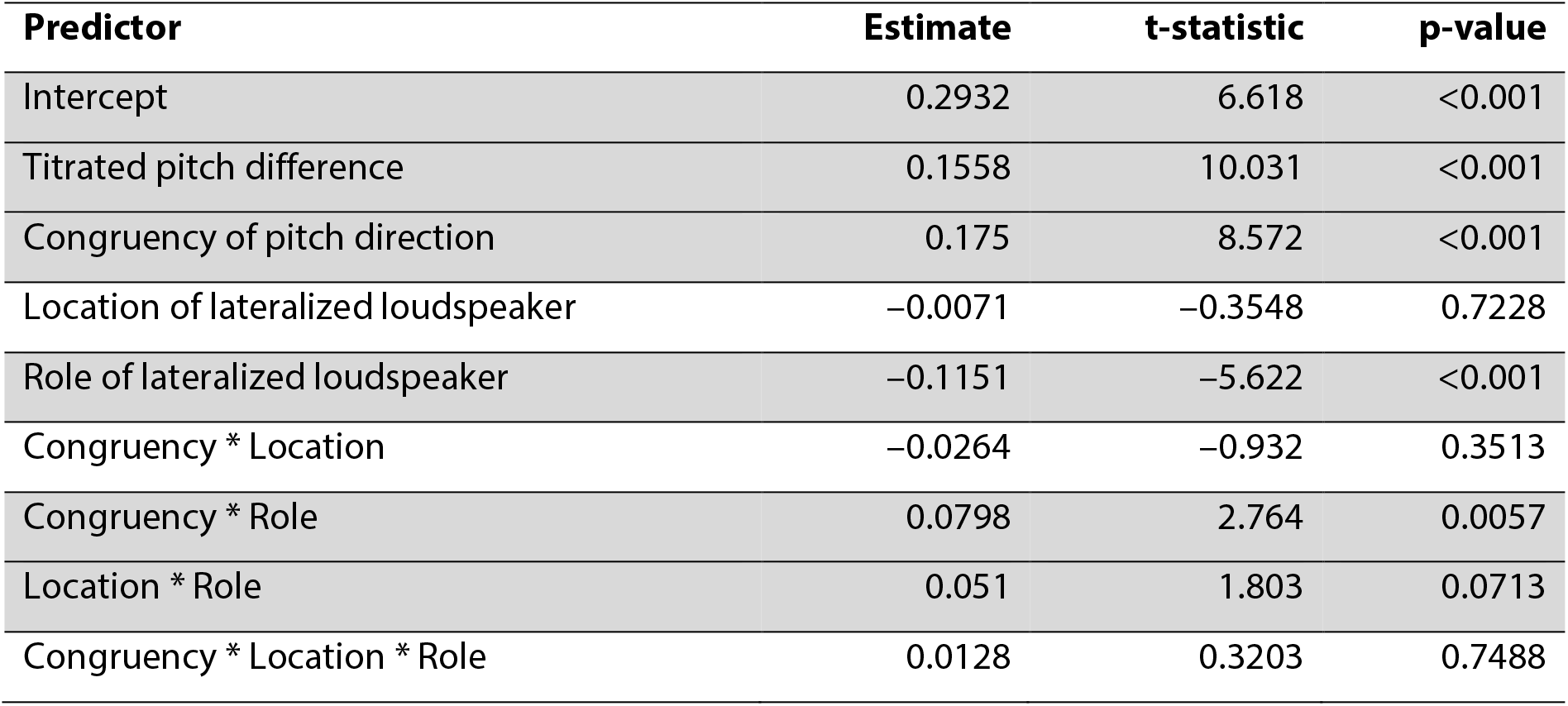
Summary table of a linear mixed-effects model to predict the outcome variable confidence-weighted accuracy, using the model formula: Confidence-weighted accuracy ~ 1 + Titrated pitch difference + Congruency of pitch direction * Location of lateralized loudspeaker * Role of lateralized loudspeaker + (1|Participant ID). Degrees of freedom = 16949. Rows for predictors with p < 0.1 have gray shading.

**Figure 2-2.**
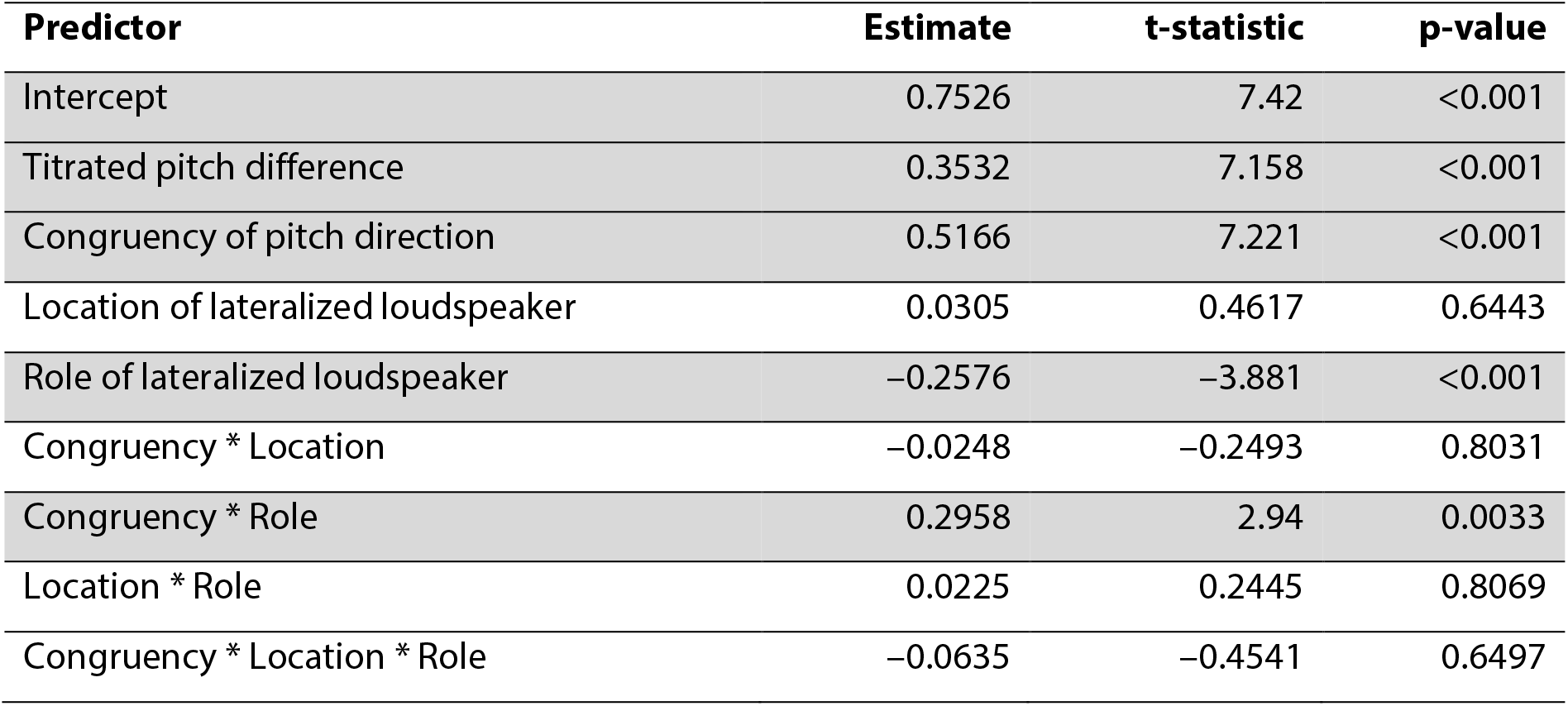
Summary table of a generalized linear mixed-effects model (logit link function) to predict the outcome variable raw accuracy, using the model formula: Raw accuracy ~ 1 + Titrated pitch difference + Congruency of pitch direction * Location of lateralized loudspeaker * Role of lateralized loudspeaker + (1|Participant ID). Degrees of freedom = 17005. Rows for predictors with p < 0.1 have gray shading.

**Figure 2-3.**
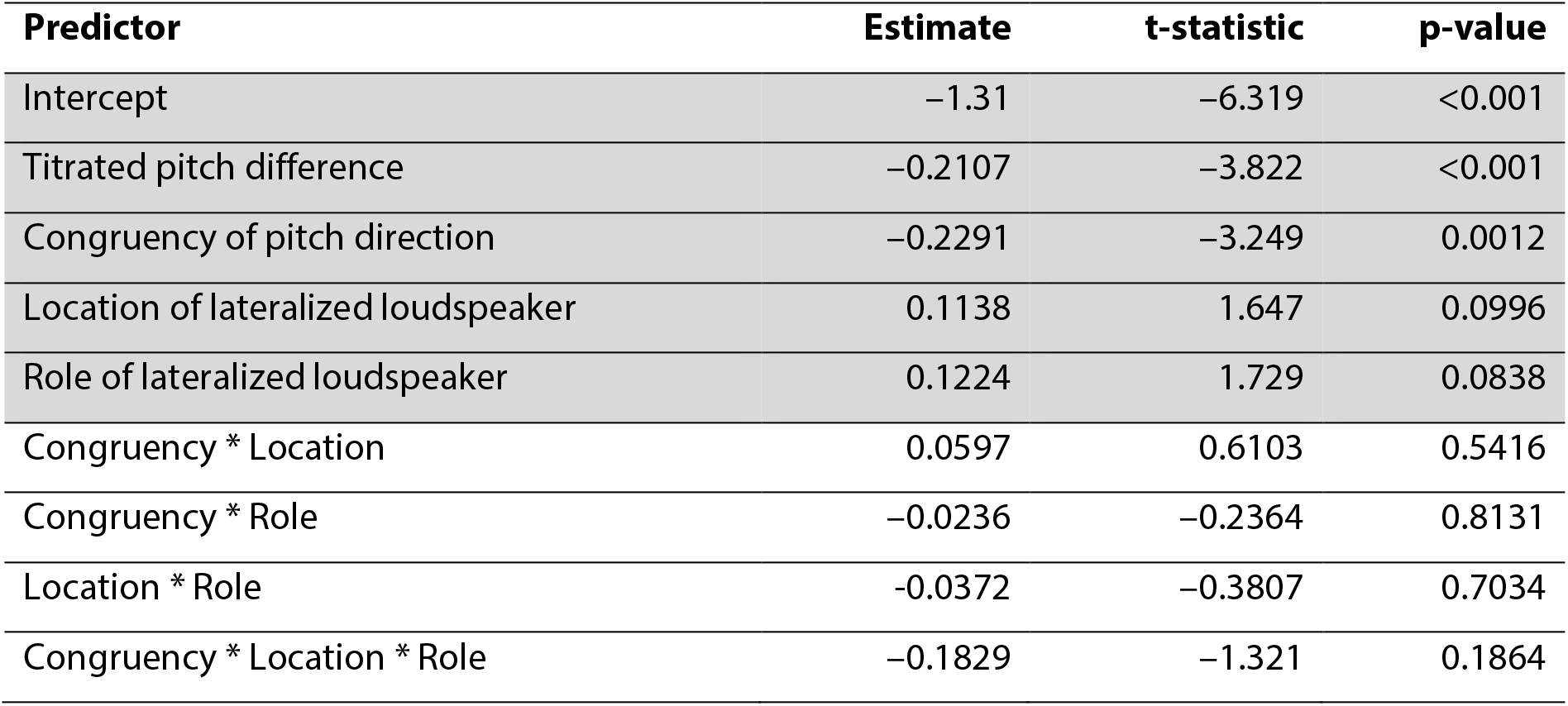
Summary table of a linear mixed-effects model to predict the outcome variable log-transformed response time, using the model formula: Log-transformed response time ~ 1 + Titrated pitch difference + Congruency of pitch direction * Location of lateralized loudspeaker * Role of lateralized loudspeaker + (1|Participant ID). Degrees of freedom = 17005. Rows for predictors with p < 0.1 have gray shading.

**Figure 3-1.**
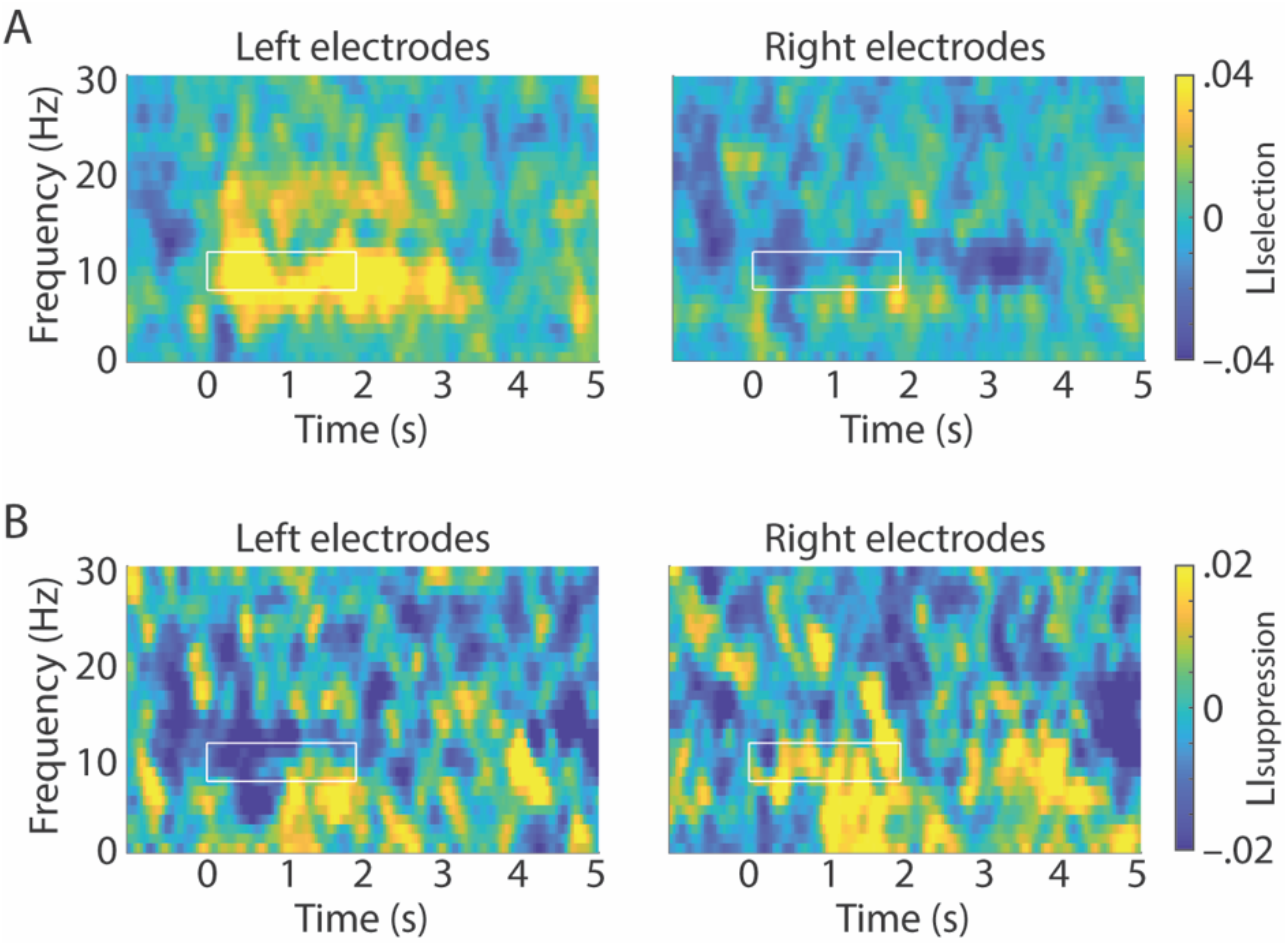
Time-frequency representations on the left and right show the respective grand-average lateralization index at 12 left- and 12 right-hemispheric electrodes (electrodes highlighted in Fig. 3 B&D). (A) Lateralization index for the selection of lateralized targets (LI_selection_). (B) Lateralization index for the suppression of lateralized distractors (LI_suppression_). White outlines indicate the time-frequency region of interest before the onset of tone sequences (0–1.9 s; 8–12 Hz).

